# Primary cilia drive postnatal tidemark patterning in articular cartilage by coordinating responses to Indian Hedgehog and mechanical load

**DOI:** 10.1101/2021.07.04.451018

**Authors:** Danielle Rux, Kimberly Helbig, Biao Han, Courtney Cortese, Eiki Koyama, Lin Han, Maurizio Pacifici

## Abstract

Articular cartilage (AC) is essential for body movement, but is highly susceptible to degenerative diseases and has poor self-repair capacity. To improve current subpar regenerative treatment, developmental mechanisms of AC should be clarified and, specifically, how postnatal multi-zone organization is acquired. Primary cilia are cell surface organelles crucial for mammalian tissue morphogenesis and while the importance of chondrocyte primary cilia is well appreciated their specific roles in postnatal AC morphogenesis remain unclear. To explore these mechanisms, we used a murine conditional loss-of-function approach (*Ift88-flox*) targeting joint-lineage progenitors (*Gdf5Cre*) and monitored postnatal knee AC development. Joint formation and growth up to juvenile stages were largely unaffected, however mature AC (aged 2 months) exhibited disorganized extracellular matrix, decreased aggrecan and collagen II due to reduced gene expression (not increased catabolism), and marked reduction of AC modulus by 30-50%. In addition, we discovered the surprising findings that tidemark patterning was severely disrupted and accompanied alterations in hedgehog signaling that were also dependent on regional load-bearing functions of AC. Interestingly, *Prg4* expression was also increased in those loaded sites. Together, our data provide evidence that primary cilia orchestrate postnatal AC morphogenesis, dictating tidemark topography, zonal matrix composition and mechanical load responses.

## INTRODUCTION

Body movement requires uniquely shaped synovial joints to maximize range and type of motion based on anatomical location and load-bearing needs (Mow & Sugalski, 2001). Much has been learned about the structure and function of adult joint tissues –articular cartilage (AC), meniscus and intra joint ligaments– including the biomechanical and lubricating roles each tissue plays in sustaining motion and resilience (Longobardi et al., 2015). While it is clear that the majority of morphogenesis and terminal organization of mature AC occurs during postnatal life in coordination with growth of the skeleton (Decker, 2017; Decker et al., 2017; Kozhemyakina et al., 2015; L. Li et al., 2017), cellular mechanisms remain poorly understood. A full understanding of this morphogenetic process would be critical to inform on how joint developmental defects/diseases are established and, as importantly, how defective and damaged joint tissues could be repaired and regenerated.

In adults, fully mature AC displays a stereotypic zonal organization of cells and extracellular matrix (ECM) that contains both non-mineralized and mineralized layers. The non-mineralized hyaline cartilage includes: (i) a top superficial zone of flat cells juxtaposed with the synovial space and parallel to the surface, producing ECM components such as lubricin/*Prg4*; (ii) an intermediate zone of round chondrocytes amidst aggrecan and randomly aligned collagen II fibrils; (iii) a thick, deep zone of polarized chondrocytes oriented into columns and containing abundant ECM rich in aggrecan and with collagen II fibrils aligned perpendicular to the surface. The mineralized cartilage comprises a bottom zone with large hypertrophic-like chondrocytes that conjoins AC to the underlying subchondral bone (Hunziker et al., 2007). The tidemark represents the mineralization front demarcating the boundary between non-mineralized and mineralized cartilage. While, little is known of is developmental origins, significant evidence exists that the tidemark advances toward the cartilage surface with age (Johnson, 1962; Lane & Bullough, 1980; Loeser et al., 2012; Oegema et al., 1997; Radin et al., 1991) or with alterations in mechanical load (Nomura et al., 2017; O’Connor, 1997; Tomiya et al., 2009), reducing non-mineralized cartilage thickness. In addition, the transitioning of aquatic axolotl salamanders to terrestrial life results in the formation of a tidemark (Cosden-Decker et al., 2012; Rux et al., 2019), suggesting ambulation influences initial positioning. Despite these observations, no cellular mechanisms on tidemark patterning or disease-state repositioning have been clarified.

Postnatal AC morphogenesis is complex as it needs to eventually establish a highly structured, anisotropic tissue critical for joint mechanical function (Gannon et al., 2015; Hunziker et al., 2007; Julkunen et al., 2009). Recent studies by our group and others have led to a better understanding of the basic tenets of postnatal morphogenesis of AC in mouse knee joints (Decker et al., 2017; Kozhemyakina et al., 2015; L. Li et al., 2017). At birth, nascent AC is disorganized and contains a pool of highly proliferative progenitors which give rise to all of the cells in adult AC. By 2-3 weeks of age, AC is matrix-rich, but still lacks a clear zonal organization. By 8 weeks, the stereotypical zonal organization is established. Importantly, we found that cell proliferation decreases rapidly soon after birth while the tissue continues to thicken through matrix production/accumulation and increases in average cell size (Decker et al., 2017). Thus, there are three stages to postnatal AC development: proliferation (neonatal), growth (adolescent) and maturation (adult). Lineage tracing in our recent study strongly suggested that chondrocytes establish zonal organization and vertical columns during the maturation phase (Decker et al., 2017), but the mechanisms by which this could occur have not been elucidated.

Primary cilia are putative mechano- and morphogen-transducing cell surface organelles present on nearly every cell in mammals and are crucial for tissue morphogenesis. Cytoskeletal regulation, cell polarity and a number of signaling pathways (e.g., hedgehog, wnt and TGFβ) are governed, at least in part, by the primary cilium (Goetz & Anderson, 2010; Wheway et al., 2018). The chondrocyte primary cilium is an important regulator of cartilage function (Haycraft et al., 2007; Serra, 2008; Yuan & Yang, 2016) and lengthening has been correlated with human osteoarthritis (OA) (McGlashan et al., 2008). In mice, conditional loss of function in AC results in deficits that often lead to early onset osteoarthritis (OA) (Chang et al., 2012; Coveney et al., 2021; Irianto et al., 2014; Sheffield et al., 2018; Song et al., 2007; Yuan & Yang, 2015). It is thus clear that primary cilia are important to AC health, but the specific mechanisms by which they function during postnatal morphogenesis remain unclearly defined.

In an effort to identify mechanisms of postnatal AC development in mice and how cilia function in this regard, we developed a joint-specific (*Gdf5Cre*) disruption of primary cilia (*IFT88-flox*) and delineated AC development up to 8 weeks of age. We found that while the disruption of primary cilia does not affect embryonic or early postnatal growth through 3 weeks of age, it significantly altered AC maturation including ECM production and organization, culminating in reduced AC modulus at 8 weeks of age. Surprisingly, we also found significant disruptions in tidemark patterning that altered the organization of mineralized and non-mineralized zones. Our mechanistic studies revealed primary deficits in hedgehog signaling starting at 3 weeks accompanied by secondary changes to BMP and TGFβ signaling by 8 weeks. Importantly, our data indicate that the hedgehog pathway drives tidemark patterning and location during postnatal AC morphogenesis, being the first to identify a signaling mechanism that is capable of regulating and altering the tidemark.

## RESULTS

### *Gdf5Cre* is joint specific and conditional deletion of *IFT88* results in loss of primary cilia

In order to target primary cilia specifically in synovial joints, we used *Gdf5Cre* in combination with an *IFT88* conditional allele. *Gdf5* is a unique marker for the embryonic mesenchymal progenitor cells of synovial joints called the interzone. *Gdf5Cre* faithfully marks these cells in the embryo and lineage-traces to all of the mature tissues of synovial joints (Decker et al., 2017; Koyama et al., 2008) (Fig. 1Aa-c) but not growth plate chondrocytes (Fig. 1Ad). In order to study primary cilia, we combined *Gdf5Cre* with an *IFT88* conditional allele whose product (Intraflagellar Transport Protein 88) is important in anterograde transport in cilia and maintenance of the cilium structure. The result is loss of primary cilia in *Gdf5*-expressing interzone cells and in adult synovial joints. We visualized primary cilia using an antibody against acetylated tubulin and found they were present on nearly every chondrocyte in control AC (Fig. 1Ba-b) and in the growth plate (Fig. 1Bc and Bf) but a significant loss in all zones of AC in mutant animals (Fig. 1Bd-e) confirming our model was reliable.

**Figure 1.**
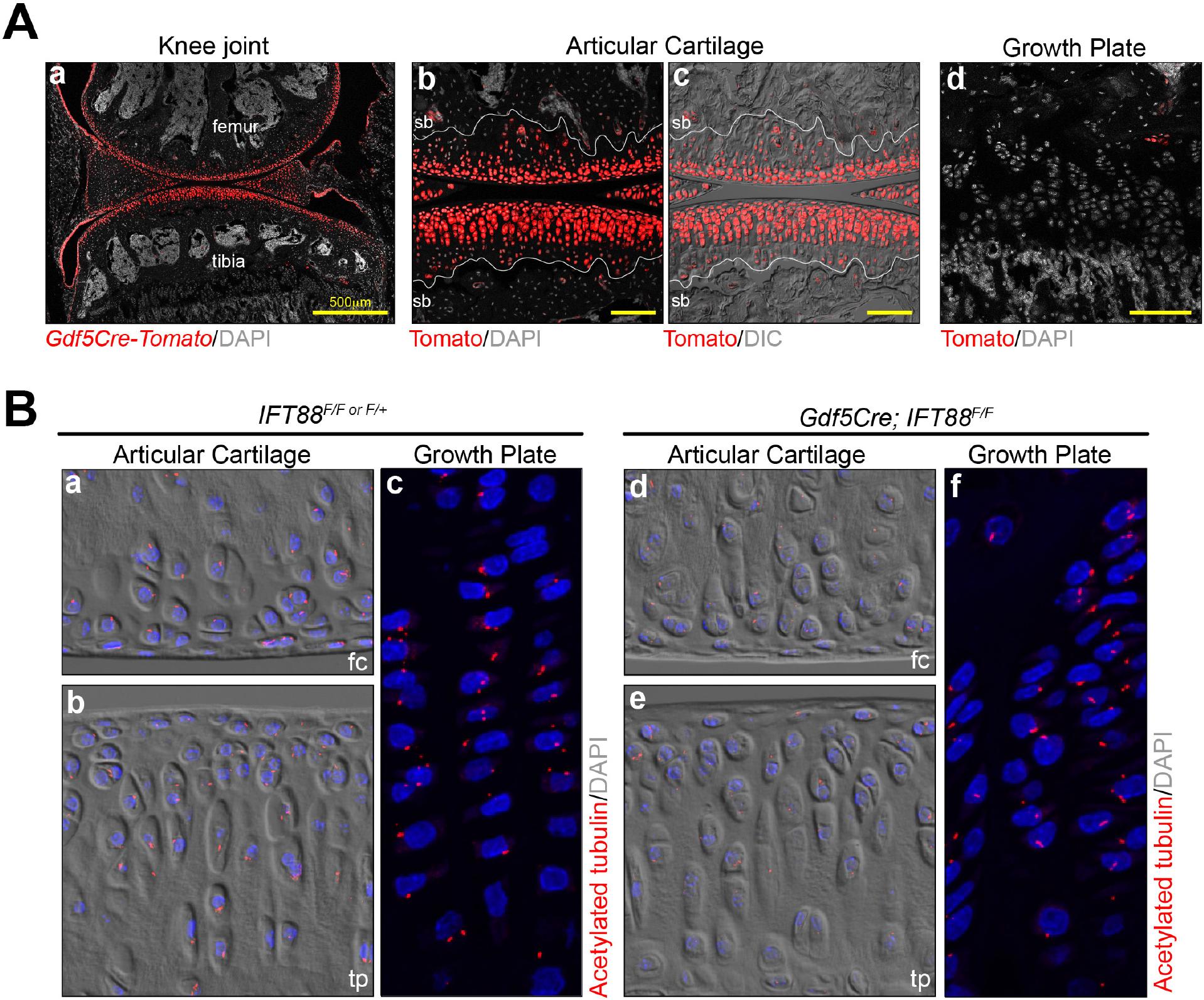
*Gdf5Cre* is joint specific and its conditional deletion of *IFT88* results in loss of primary cilia. (**A**) *Gdf5Cre/+; Rosa-tdTomato* lineage cells are present in adult joints (a) and all zones of AC as shown with DIC imaging to visualize the tidemark (b-c), but are absent in growth plate (d). (**B**) Primary cilia are visualized by acetylated tubulin staining. They are present in control AC (a-b) and growth plate chondrocytes (c); in mutants, they are absent from AC (d-e) but still present in growth plate chondrocytes (f). sb, subchondral bone; fc, femoral condyle; tp, tibial plateau. scale bar = 100 µm unless otherwise noted.

### Joint-specific deletion of *IFT88* leads to disrupted AC tidemark patterning and inferior biomechanical function

Next, we evaluated histology and performed mechanical testing to assess the consequences of loss of primary cilia during postnatal AC morphogenesis. We used safranin O staining to visualize basic histology and proteoglycans in the ECM (Fig. 2A). At birth, nascent AC of mutants was indistinguishable from controls (Fig. 2Aa-b, brackets). Up to 3 weeks, immature AC developed similarly (Fig. 2Ac-d) and higher magnification images revealed no discernible differences in cell morphology or patterning (Fig. 2Ae-h). However, at 8 weeks we noted irregular accumulation of the interterritorial ECM (arrows) and chondrocyte size (asterisks) in the femoral condyle (Fig. 2Ai-l) and a consistent reduction of safranin O in the tibial plateau (Fig. 2i-j, m-n). Thus, the loss of IFT88 primarily affected the maturation phase of postnatal AC morphogenesis. Growth plates were unaffected at all stages consistent with no recombination from *Gdf5Cre* (Fig. S1). Next, we utilized a combination of safranin O-stained sections and sections imaged with Differential Interference Contrast (DIC) to quantify AC thickness, AC composition of mineralized and non-mineralized cartilage and also cell density. These analyses revealed several notable changes. First, in healthy AC, the tidemark (Fig. 2B, dashed yellow line) closely followed the contour of the femoral condyle surface 2-3 cell layers below, but was relatively flat in the load-bearing region of the tibial plateau located about 5-6 cell layers from the surface (Fig. 2Ba, bracket). In mutants, the tidemark was often difficult to visualize, but followed an irregular path in the femoral condyle and was flat, but noticeably further from the surface in the tibial plateau (Fig. 2Bb, bracket). Quantifying AC composition as a percent area, we found that non-mineralized cartilage constituted approximately 31% of the femoral condyle and 51% of tibial plateau AC in control animals; this increased slightly in mutant femoral condyle AC to 36% and significantly in tibial plateau AC to 70% (Fig. 2C). For consistency, we measured AC height at its thickest regions in each tissue section. We found no changes in AC thickness at 3 weeks nor in tibial plateau AC at 8 weeks, but the AC height was significantly higher in the thickest regions of mutant femoral condyles (Fig. 2Da-b). Bisected into mineralized and non-mineralized cartilage at 8 weeks, we found similar trends in composition from Fig. 2C. In the femoral condyle, mutants showed a significant increase in non-mineralized thickness, but no change mineralized thickness (Fig. 2Dc) and in the tibial plateau, mutants showed a significant increase in non-mineralized cartilage and decrease in mineralized cartilage (Fig. 2Dd). Thus, AC composition in tibial plateau is consistently more non-mineralized whereas mineralization is much more variable in the femoral condyle but follows the same trend. Interestingly, we found that cell density in non-mineralized AC was significantly reduced in both regions of mutant AC (Fig. 2E).

**Figure 2.**
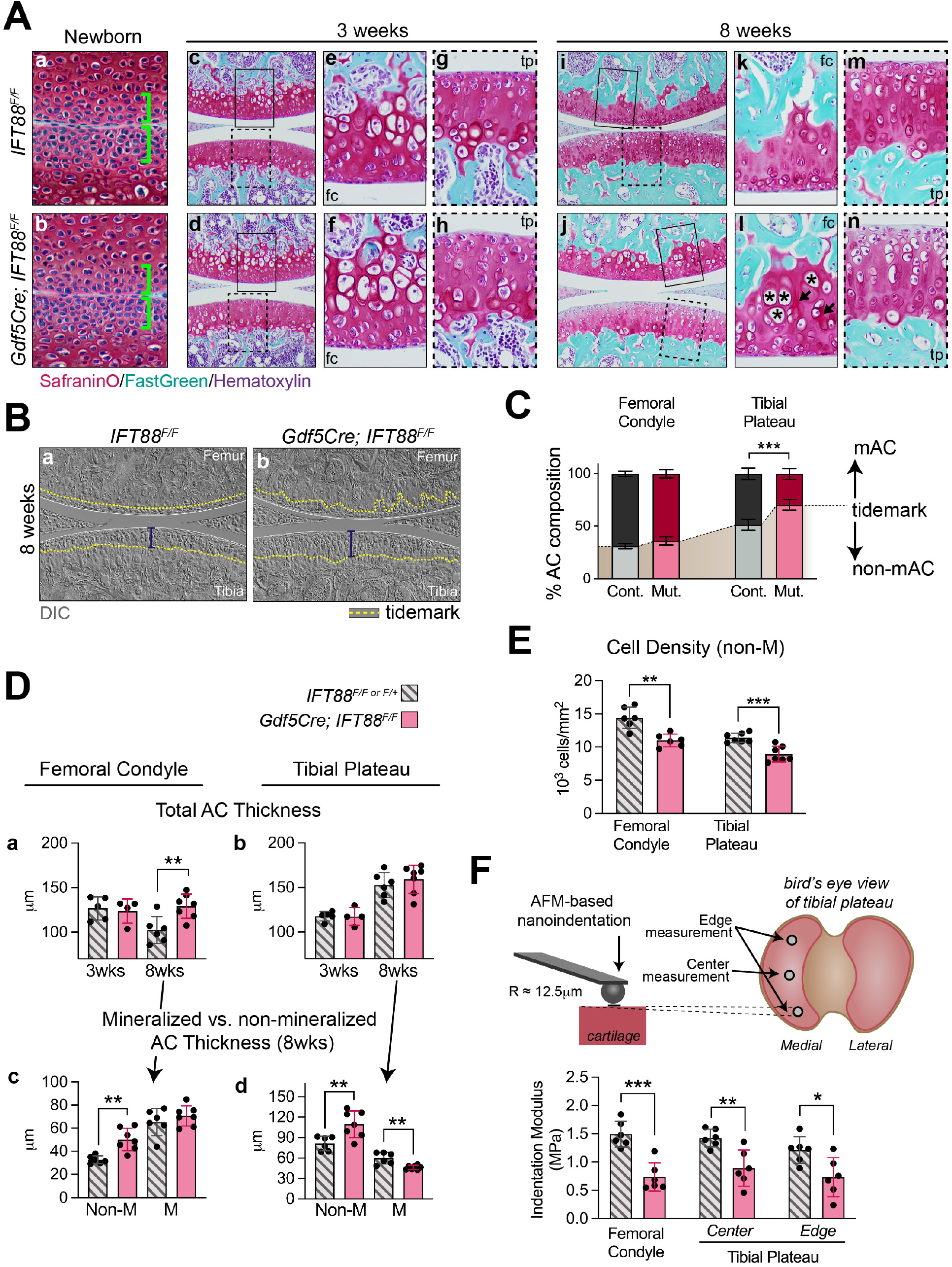
Joint-specific deletion of *IFT88* results in disrupted AC tidemark patterning and inferior biomechanical function. (**A**) Safranin O staining at birth through 8 weeks. Nascent AC at birth is unchanged in mutant joints (a-b, brackets). Low magnification (c-d) and high magnification images of femoral condyle (e-f) or tibial plateau (g-h) are comparable in controls and mutants at 3 weeks. At 8 weeks, control AC (i, k, m) displays strong safranin O staining throughout the femoral condyle (k) and tibial plateau (m). In mutant AC (j, l, n), the femoral condyle displays aberrant ECM deposition (l, arrows) and increased cell size (l, asterisks), while the tibial plateau displays reduced Safranin O staining (n). (**B**) DIC imaging of the tidemark (dashed yellow line) reveals ordered patterning in control AC (a). In mutant AC, the tidemark of the femoral condyle is highly disorganized and its distance from the surface is increased in the tibial plateau (b, bracket). (**C**) Mineralized (mAC) and non-mineralized (non-mAC) composition is shown as a percent area from multiple tissue sections and is significantly altered in mutant tibial plateau. (**D**) Measurements of AC height at the thickest point among tissue sections reveals a significant increase in mutant femoral condyle at 8 weeks (a) but no change in tibial plateau (b). Bisected into non-mineralized (non-M) and mineralized (M) thickness (c-d), mutants display a thicker non-mineralized zone in both locations, and a reduced mineralized zone in the tibial plateau. (**E**) Cell density in the non-mineralized AC is decreased in all knee joint AC. (**F**) Biomechanical function was measured by AFM-nanoindentation from 3 regions. Schematic shows approximate location of the 2 regions of tibial plateau AC. Indentation modulus was significantly decreased in all locations. fc, femoral condyle; tp, tibial plateau scale bar = 100 µm.

To assess possible functional consequences of these morphologic changes in mature AC, we used atomic force microscopy (AFM)-nanoindentation to measure AC stiffness. We measured two regions on the medial tibial plateau (the edge covered by meniscus and the center load-bearing region) and one region on the femoral condyle. In all regions of interest, the indentation modulus (i.e. resistance to deformation) was significantly decreased in mutants (Fig. 2F). Thus, loss of IFT88 in AC results in impaired load-bearing function during tissue maturation.

### IFT88 regulates chondrocyte fate and matrix deposition in mineralized and non-mineralized zones

Next, we performed immunofluorescence staining for several key ECM and chondrocyte-specific markers. Collagen II is a major structural component of cartilage ECM, responsible for tensile strength of the tissue (Mow & Ratcliffe, 1997). In control animals, it was uniformly dispersed in AC at 3 weeks and 8 weeks (Fig. 3Aa and Ac). In mutant animals, we noted no changes at 3 weeks (Fig. 3Ab) and a mild reduction at 8 weeks (Fig. 3Ad). Collagen X is a marker of hypertrophic and mineralizing chondrocytes. We found collagen X similarly localized to cells juxtaposed to the developing secondary ossification center in control (Fig, 3Ba, c) and in mutant AC (Fig. 3Bb, d) consistent with a lack of precocious hypertrophy closer to the AC surface. Aggrecan is the major proteoglycan in cartilage ECM that gives it its compressive load-bearing and energy dissipation properties. In healthy tissue, it was uniformly deposited in the entire thickness of developing AC at 3 weeks (Fig. 3Ca) and was enriched in non-mineralized cartilage at 8 weeks (Fig. 3Cc). A clear boundary (i.e., tidemark) could be visualized where the aggrecan-low mineralized cartilage met the aggrecan-rich non-mineralized cartilage (Fig. 3Cc, arrows). In IFT88 mutants, aggrecan deposition was comparable to controls in immature cartilage at 3 weeks (Fig. 3Cb), but notable disorganization, lack of tidemark and variable deposition (high, arrows and low, arrowheads) were all apparent in mature cartilage at 8 weeks (Fig. 3Cd). Sox9 is considered a master transcriptional regulator of chondrocyte fate and can directly regulate the expression of aggrecan and collagen II (Lefebvre et al., 2019). In immature AC, Sox9 was present uniformly in articular chondrocytes of controls and mutants at 3 weeks (Fig. 3Da-b) consistent with the uniform distribution of collagen II and aggrecan at this stage. In mature AC, Sox9-positive cells were largely restricted to the non-mineralized cartilage (Fig. 3Dc), as visualized by DIC imaging (Fig. S2a-c), and is consistent with a recent report of Sox9 function in AC (Haseeb et al., 2021). Mutant tissues at 8 weeks continued to express Sox9 broadly in non-mineralized chondrocytes (Fig. S2d-f), but Sox9-positive cells could also be seen in articular chondrocytes beneath the tidemark in the femoral condyle (Fig. 3Dd, arrows). Taken together, these data show that IFT88 mutants display an altered pattern of non-mineralized chondrocytes and coordinately disrupted ECM in mature AC, but not immature AC.

**Figure 3.**
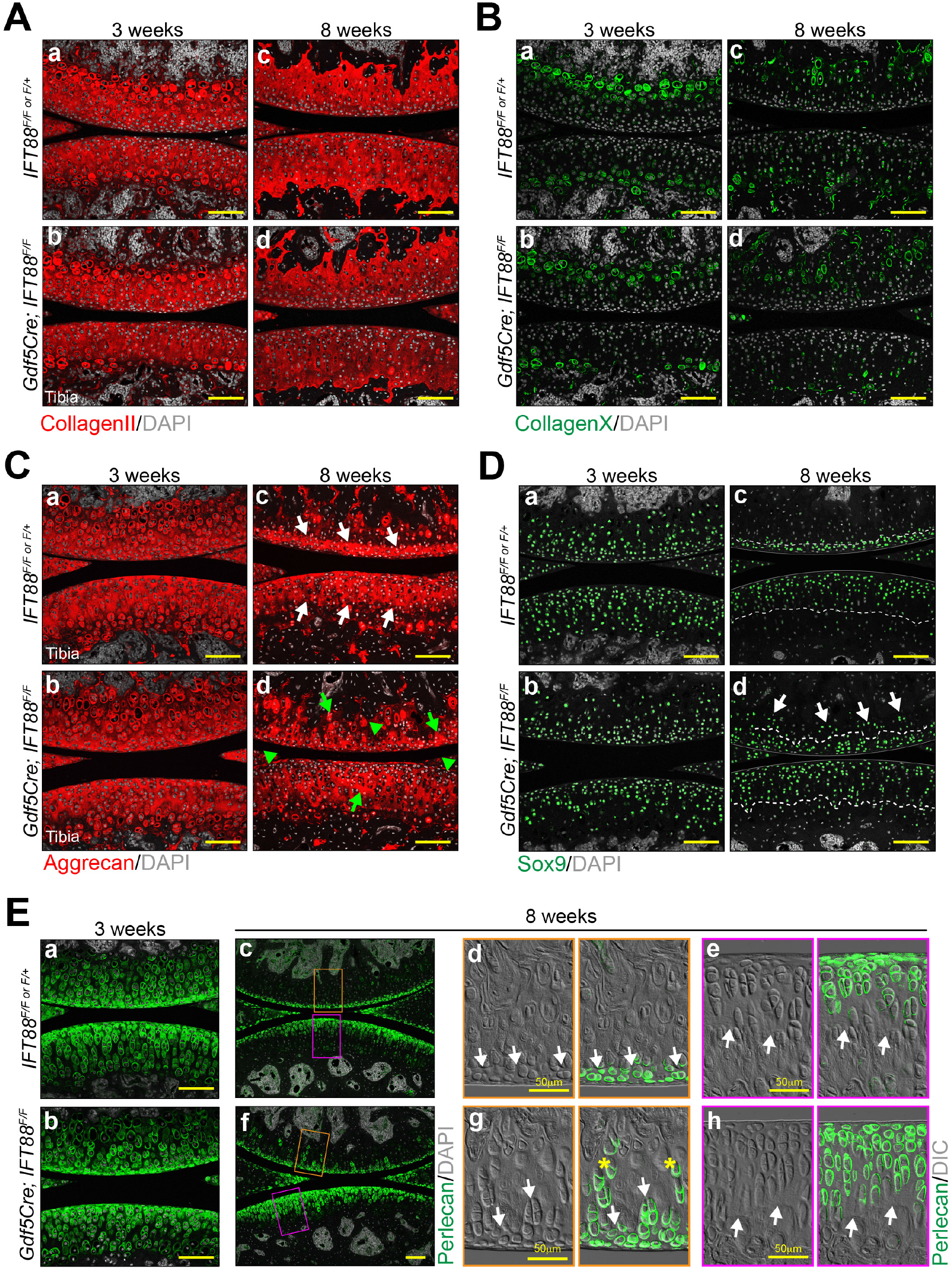
IFT88 regulates chondrocyte fate and matrix deposition in mineralized and non-mineralized zones. (**A**) Collagen II immunostaining is broadly distributed in AC and comparable at 3 weeks (a-b) and moderately reduced at 8 weeks in mutants (c-d). (**B**) Collagen X immunostaining is restricted to chondrocytes near the subchondral bone and is comparable at 3 weeks (a-b) and 8 weeks in all samples (c-d). (**C**) Aggrecan immunostaining is broadly distributed and comparable at 3 weeks (a-b). It is enriched in non-mineralized 8 week AC of controls (c, white arrows point to tidemark) and is disorganized in 8 week AC of mutants (d), with random areas of high staining (green arrows) and low staining (green arrow heads). (**D**) Sox 9 is broadly distributed and comparable in AC at 3 weeks (a-b). It is also broadly distributed in non-mineralized AC of controls and mutants at 8 weeks (c-d, dashed line represents the tidemark), but mutant femoral condyle AC also displays ectopic Sox9-positive cells beyond the tidemark (d, white arrows). (**E**) Perlecan is broadly distributed in AC and comparable at 3 weeks (a-b) and is enriched nearest the surface at 8 weeks (c, f). High magnification with DIC shows the tidemark (d, e, g, h, white arrows). Perlecan+ cells are present below the tidemark in mutant femoral condyle (g, asterisks) and in deeper cell layers of mutant tibial plateau (h). scale bar = 100 µm.

Pericellular matrix (PCM) creates a thin microenvironment surrounding chondrocytes within the broader ECM and is considered a crucial transducer of biomechanical signals (Guilak et al., 2006). To observe any changes to the PCM, we performed immunofluorescence staining for perlecan, a biomarker of cartilage PCM (Kvist et al., 2008; Wilusz et al., 2012). In immature AC at 3 weeks, perlecan was broadly distributed in both control and mutant tissues (Fig. 3Ea-b). In mature AC 8 weeks, perlecan was localized to chondrocytes nearest to the surface in control tissue (Fig. 3Ec) and high magnification images overlaid with DIC imaging revealed restriction to the non-mineralized cartilage above the tidemark (Fig. 3Ed-e, arrows). The organization of perlecan deposition was significantly disrupted in mutant AC at 8 weeks (Fig. 3Ed). In the femoral condyle, chondrocytes with perlecan-rich PCM could be visualized far below the tidemark (Fig. 3Eg, asterisks), and in tibial plateau AC there was an expansion of perlecan-positive chondrocytes in the load-bearing region above the tidemark (Fig. 3Eh). Together, these data provided strong evidence that loss of IFT88 in AC disrupts tidemark patterning and chondrocyte fate/microenvironment during postnatal maturation.

### Loss of IFT88 in AC does not increase matrix degradation

Previous *in vitro* studies have shown that primary cilia, and IFT proteins, regulate ECM proteases in chondrocytes (Coveney et al., 2018). To test whether this applied to our *in vivo* model, we performed immunofluorescence staining or *in situ* hybridization for MMP13 (collagenase), Adamts4 and Adamts5 (aggrecanases) at 8 weeks. MMP13 was unchanged in mutants and was restricted to the base of AC at the transition to subchondral bone (Fig. 4A). We used RNAscope (FastRed detection) to perform *in situ* hybridization and took advantage of FastRed fluorescence for imaging low-expressing genes. In AC, we found no expression of either *adamts4* or *adamts5* in control or mutant AC at 8 weeks (Fig. 4B-C). To ensure these data were reliable, we compared to other tissues in and around the joint. *Adamts4* (Fig. S3A) was expressed in primary spongiosa beneath the growth plate (PS, arrowheads) and in periosteum (PO, arrows) and *Adamts5* (Fig. S3B) was highly expressed by inner synovial fibroblasts in both controls and mutants. These data indicate that the changes observed in aggrecan and collagen II deposition in mutant AC (Fig. 2) were not caused by matrix degradation.

**Figure 4.**
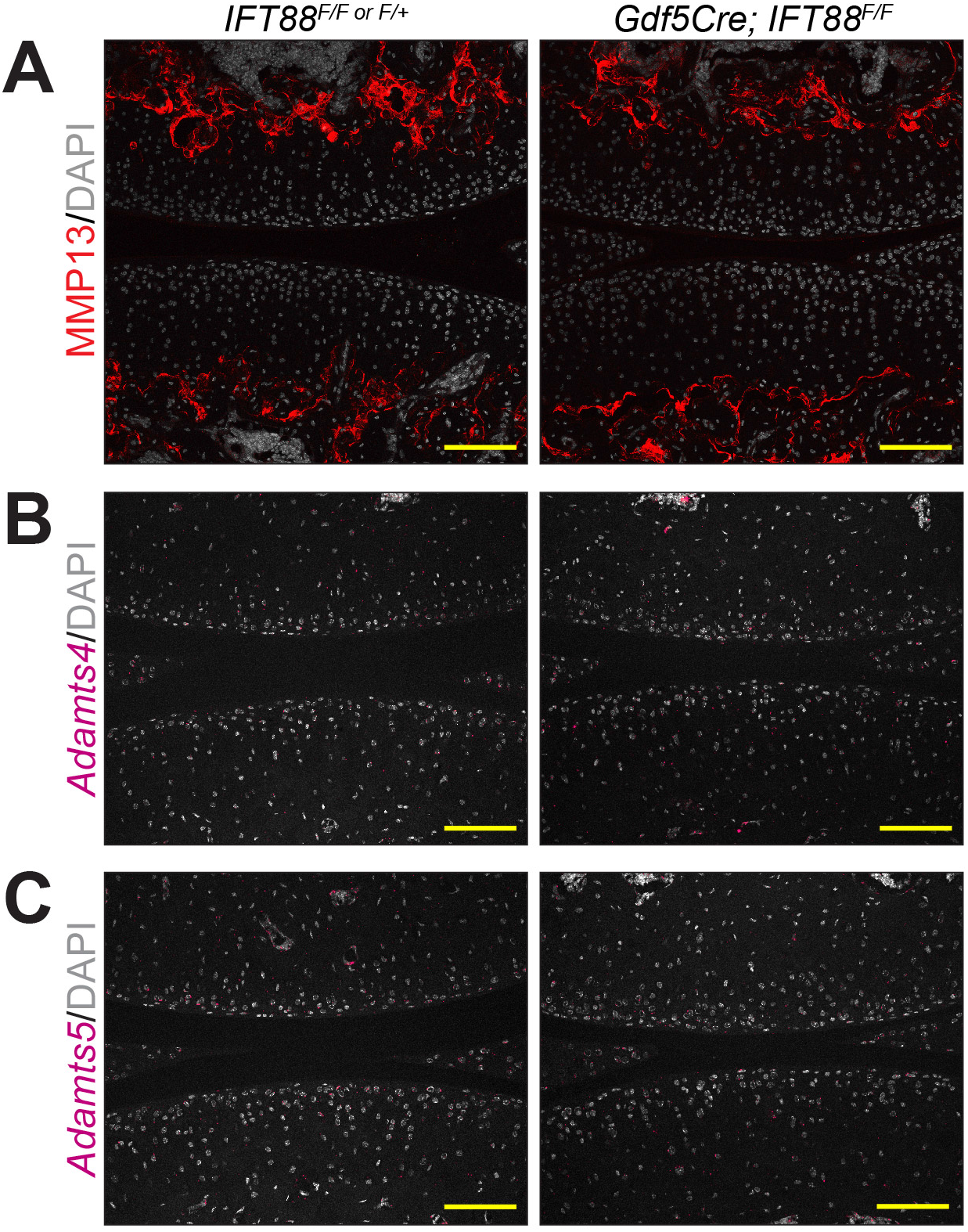
Loss of IFT88 in AC does not increase degradation at 8 weeks. (**A**) Immunofluorescence for MMP13 is unchanged in controls and mutants and is always restricted to subchondral bone. (**B-C**) *Adamts4* and *Adamts5* are not detectable in control or mutant AC. scale bar = 100 µm.

### Altered ECM in mature mutant AC are the result of transcriptional changes

To assess transcriptional changes of *Col2a1*, *Acan* and *Prg4*, we utilized RNAscope (fastRed detection) imaged these high-expressing genes with brightfield. At 3 weeks, C*ol2a1* and *Acan* were broadly expressed in immature AC of both mutants and controls (Fig. 5. Aa-d) consistent with our findings from immunofluorescent staining at this stage described above (Fig. 3A and C). *Prg4* (lubricin) is important for protecting AC from OA (Chavez et al., 2019; Coles et al., 2010; Marcelino et al., 1999; Rhee et al., 2005; Ruan et al., 2013). At 3 weeks, *Prg4* was highly expressed in cells near the surface in control animals (Fig. 5Ae) and we found a significant reduction in mutant AC (Fig. 5Af). In mature AC of control animals, *Col2a1* (Fig. 5Ba-b) and *Acan* (Fig. 5Be-f) transcripts were localized to layers of AC corresponding with the domain of non-mineralized cartilage 2-3 layers deep in the femoral condyle and 5-6 layers deep in the tibial plateau (Fig. 2Ba). In mutant AC, both *Col2a1* and *Acan* expression appeared much more disorganized particularly in the femoral condyle (Fig. 5Bc and Bg, arrows). Higher magnification images of mutants (Fig. 5Bd and Bh) and controls (Fig. 5Bb and Bf), highlighted the clear disorganization of gene expression in all mutant cartilage, consistent with dysregulated tidemark patterning, ECM deposition and deviant Sox9-positive cells outside the non-mineralized cartilage (Fig. 2 and 3). In addition, mutants displayed broadly reduced expression per cell of *Col2a1* and *Acan* in the load-bearing region of the tibial plateau compared to this region in controls. Lastly, we found that *Prg4* at 8 weeks was highly expressed and uniformly distributed in cell layers closest to the AC surface (extending a bit deeper in the tibial plateau than in the femoral condyle) (Fig. 5Bi-j, bracket). In mutants, *Prg4* in the femoral condyle displayed a patchy but overall decreased expression profile (Fig. 5Bl-k, arrows) and in the tibial plateau we found a striking increase in *Prg4* in the load-bearing region (Fig. 5Bl-k, asterisk and bracket). These data suggest that reduction/disorganization of collagen II and aggrecan deposition in mature cartilage were due to changes at the transcriptional level and that, surprisingly, *Prg4* expression was altered in both immature and mature cartilage of mutants.

**Figure 5.**
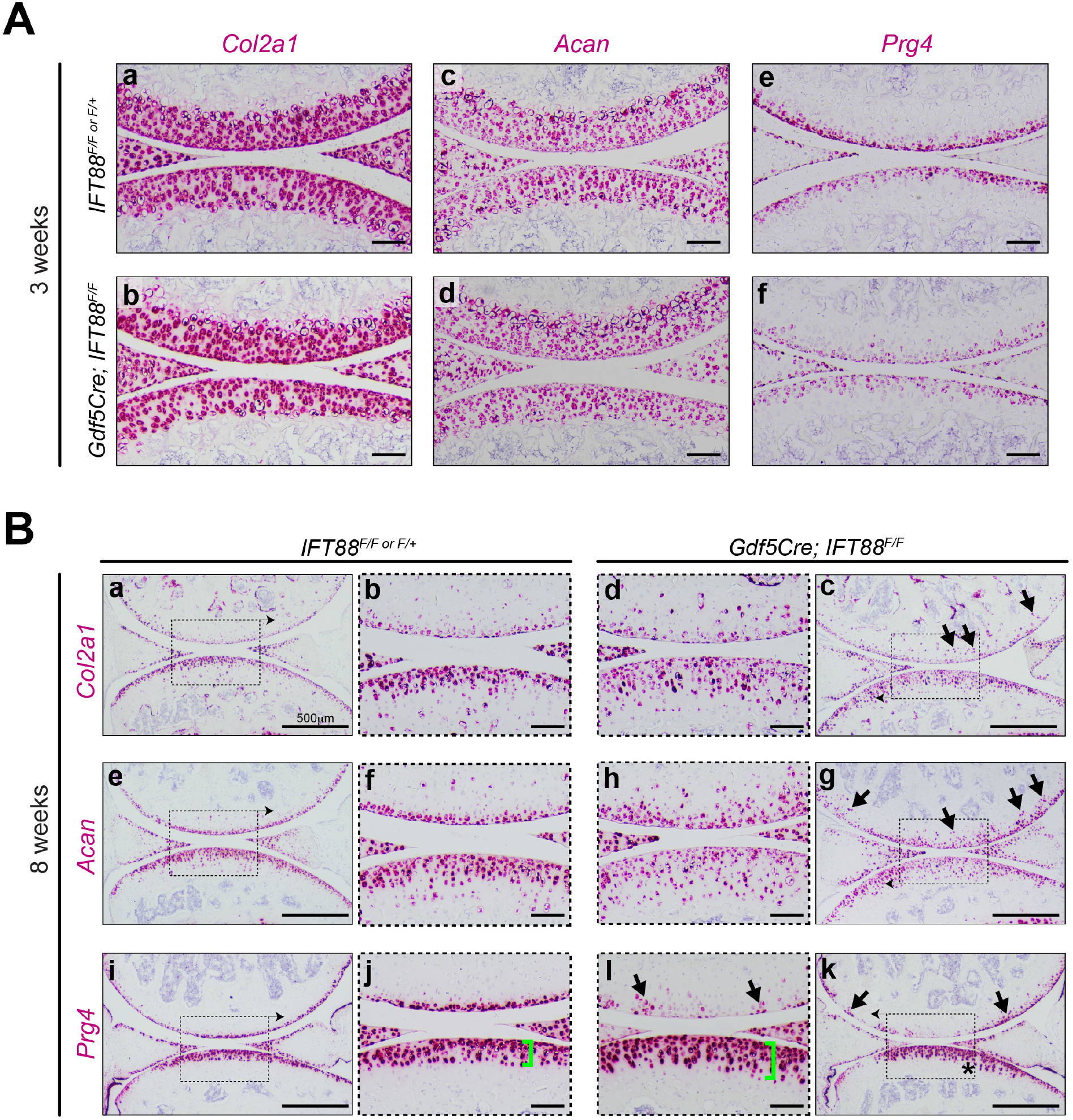
Altered ECM in mature mutant AC are the result of transcriptional changes. (a) RNAscope at 3 weeks shows *Col2a1* (a-b) and *Acan* (c-d) are comparable in controls and mutants. *Prg4* at 3 weeks is expressed close to the surface in control AC (e) and is broadly reduced in mutant tissues (f). (**B**) RNAscope at 8 weeks. Low magnification images of *Col2a1* and *Acan* (e) show organization of control AC and high magnification images (b, f) depict high levels of expression of both genes. Low magnification images of *Col2a1* (c) and *Acan* (g) show disorganization in mutant AC, particularly in the femoral condyle (arrows). High magnification images (d, h) highlight similar levels of expression to controls (b,f) in femoral condyle but decreased expression in tibial plateau AC. *Prg4* in control AC is highly expressed near the surface (i-j). *Prg4* expression in mutant AC (k-l) is decreased and patchy in the femoral condyle (arrows), while it is increased and expressed more broadly in mutant tibial plateau compared to control (asterisk and bracket). scale bar = 100 µm.

### Primary cilia impart load/location-dependent functions for key chondrogenic pathways

Primary cilia are considered a signaling nexus that have some regulatory function in most major signaling pathways (Anvarian et al., 2019). TGFβ (transforming growth factor β) and BMP (bone morphogenetic protein) regulate chondrocyte fate and ECM anabolism by distinct mechanisms via downstream SMAD-dependent mechanisms (Thielen et al., 2019). While TGFβ largely maintains non-mineralized chondrocyte function and prevents hypertrophy (Chen et al., 2012; Ferguson et al., 2000; Ionescu et al., 2003; Kang et al., 2005; Kim et al., 2012; Leboy et al., 2001; T.-F. Li et al., 2010; Seo & Serra, 2007; Yang et al., 2001), BMP can influence both non-mineralized and mineralized functions dependent on context (Bae et al., 2007; Fujii et al., 1999; Horiki et al., 2004; Javed et al., 2008, 2009; Phimphilai et al., 2006). To assess the involvement of these pathways, we used immunofluorescence for pSmad2 (TGFβ) or pSmad159 (BMP). In immature AC at 3 weeks, pSmad2-positive cells were broadly distributed in all layers (Fig. 6Aa) while pSmad159-positive cells were largely present in the deepest layers of cartilage (Fig. 6Ba). These patterns were unchanged in mutant cartilage (Fig. 6Ab and Bb). In mature AC, pSmad2 overlapped significantly with Sox9 in controls and mutants (Fig. 6C) indicating restriction to non-mineralized cartilage. The percentage of pSmad2-positive cells in control and mutant non-mineralized cartilage was unchanged but the density of pSmad2-positive cells decreased significantly in mutant tibial plateau (Fig. 6D). We performed parallel experiments for pSmad159 and found similar overlap with Sox9 in control AC and in the mutant femoral condyle, but a significant loss in mutant tibial plateau (Fig. 6E). Quantification showed significant reductions in the percent pSmad159-positive cells and also cell density in this region (Fig. 6F). These data suggest that disruption of IFT88 leads to alterations in BMP and, to a lesser degree, TGFβ signaling in the load-bearing region of the tibial plateau simultaneously with ECM disruptions already described.

**Figure 6.**
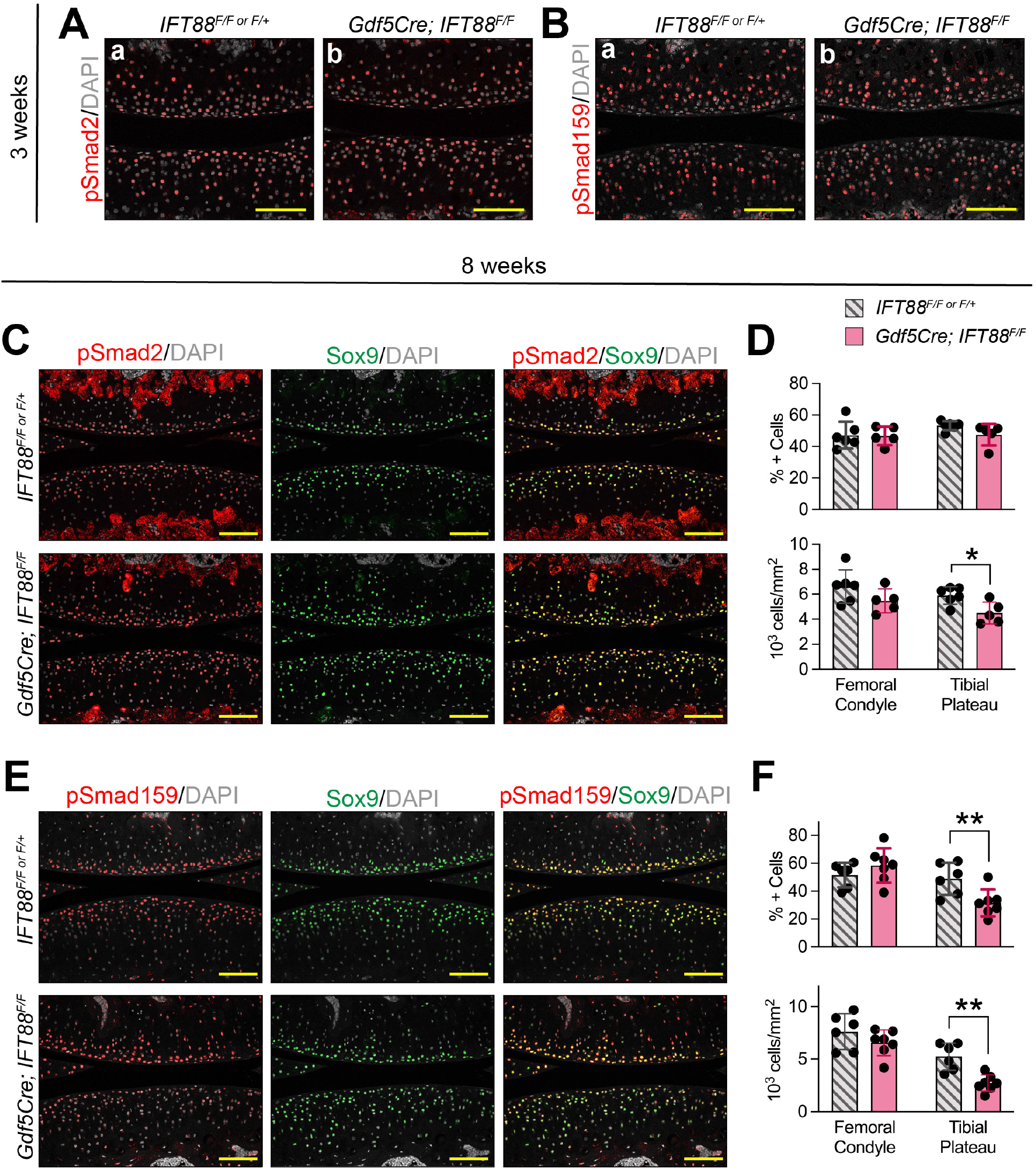
Loss of IFT88 alters TGFβ and BMP signaling dependent on anatomic location. (A) pSmad2 (TGFβ)-positive chondrocytes at 3 weeks are distributed throughout AC in both controls (a) and mutants (b). (**B**) pSmad159 (BMP)-positive chondrocytes at 3 weeks are localized in middle and bottom layers of AC in both controls (a) and mutants (b). (**C**) pSmad2 at 8 weeks overlaps significantly with Sox9 in both controls and mutants. (**D**) Quantification of non-mineralized AC shows no change in % pSmad2-positive chondrocytes in either femoral condyle or tibial plateau (top), but decreased pSmad2 chondrocyte density in the tibial plateau of mutant AC (bottom). (**E**) pSmad159 at 8 weeks overlaps significantly with Sox9 in control AC and femoral condyle of mutant AC, but not in the tibial plateau. (**F**) Quantification of non-mineralized AC shows a significant decrease in % pSmad159-positive chondrocytes of tibial plateau AC in mutants (top) and also in pSmad159 cell density in the same location (bottom). scale bar = 100 µm.

Hedgehog signaling is a major regulator of chondrocyte function and is dependent on the primary cilium for proper transduction through processing of Gli transcriptional effectors for pathway activation or repression (Wheway et al., 2018). In the absence of hedgehog ligands (Sonic, Desert or Indian), the transmembrane receptor Patched1 (Ptch1) inhibits the transmembrane protein Smoothened (Smo) to sequester Gli factors in the cytoplasm. When Hedgehog ligand is present, it binds Ptch1, the inhibition on Smo is relieved and Gli proteins are processed into active forms that translocate to the nucleus for transcriptional activation. Thus, loss of primary cilia generally results in loss of hedgehog signaling.

We sought to elucidate a possible role for hedgehog signaling in our AC model. To visualize active signaling, we used the *Gli1-LacZ* reporter mouse in combination with whole mount staining and imaging. Background staining was monitored using *Gli1-LacZ* negative tissues (Fig. S4). We noted several key findings. First, hedgehog signaling was highly active in immature AC of control animals at 3 weeks (Fig. 7A-B), and tissue sections revealed differences in *Gli1-LacZ* patterning of the femoral condyle and loaded region of the tibial plateau, localized to the first 3-4 cells layers of the femoral condyle (Fig. 7Aa) and to the bottom 3-4 cells layers in the tibial plateau (Fig. 7Bb). *Gli1-LacZ* persisted in mature AC at 8 weeks (Fig. 7E and F) with similar cell layer patterning of immature cartilage (Fig. 7Ee and Ff). *Gli1-LacZ* of immature mutant cartilage was strongly down-regulated in both the femoral condyle (Fig. 7C and Cc) and the tibial plateau (Fig. 7D and Dd). In mature mutant femoral condyle AC, *Gli1-LacZ* repression was maintained from 3 weeks (Fig. 7G and Gg) but AC in the load-bearing region of the tibial plateau exhibited very strong *Gli1-LacZ* up-regulation (Fig. 7H) including the surface zone cells that control animals did not display (Fig. 7Hh). Taken together, the data show significant hedgehog signal disruption before a structural AC phenotype is apparent, largely consistent with known roles for primary cilia in hedgehog signaling. However, our data also shows hyperactive *Gli1-LacZ* activity in the absence of primary cilia but in the presence of high mechanical load during ambulation suggesting load-dependent alterations.

**Figure 7.**
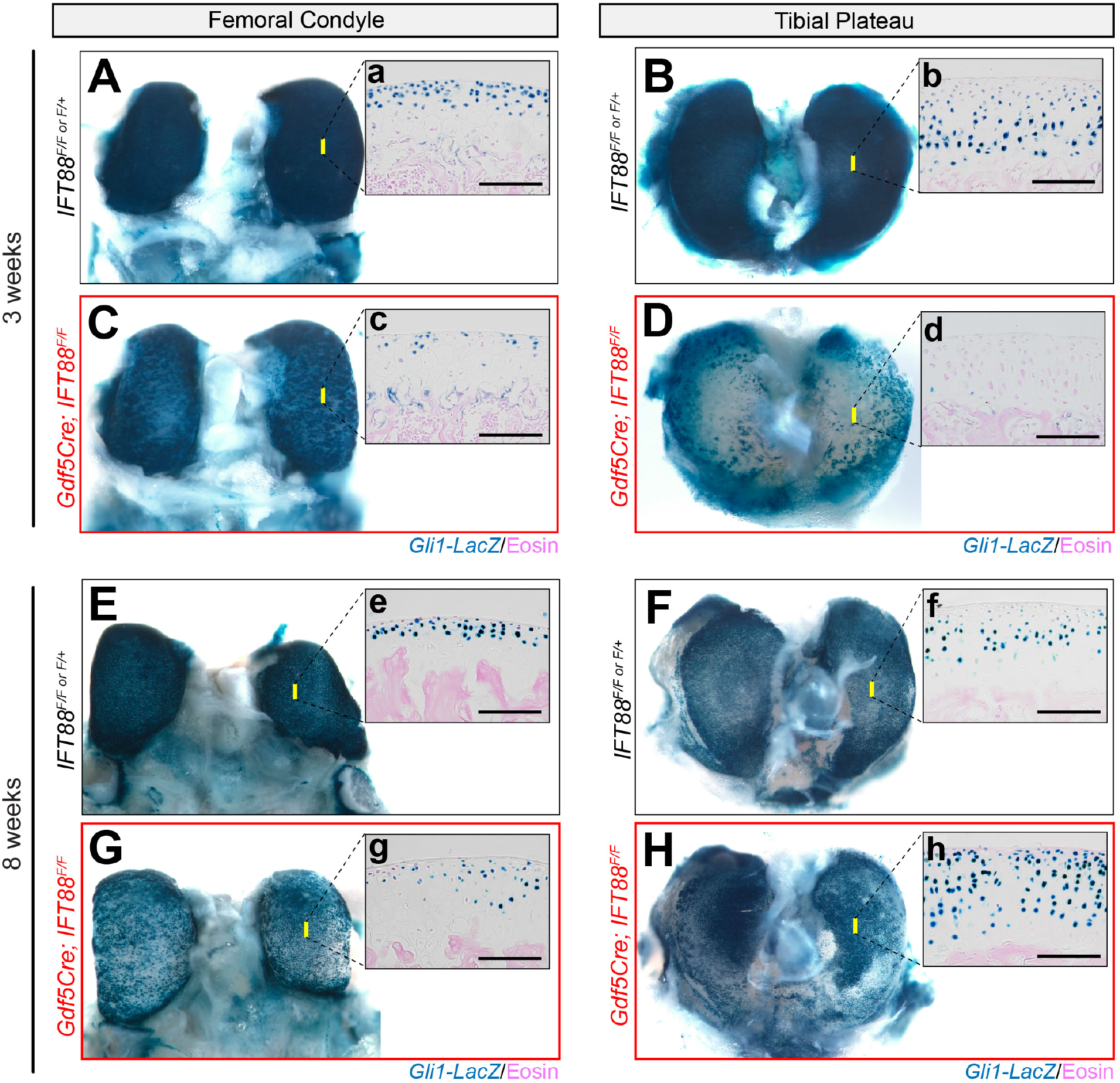
Primary cilia impart age- and location-dependent functions for Hedgehog signaling. *Gli1-LacZ* (active hedgehog signaling) was developed by whole mount β-gal staining. Yellow bars on whole mount images indicate the approximate location of tissue section images. (**A-D**) *Gli1-LacZ* at 3 weeks shows many positive cells in control AC of the femoral condyle (A) and tibial plateau (B). Tissue sections reveal that *Gli1-LacZ*-expressing cells localized near the surface in the femoral condyle AC (a) and near the subchondral bone in the tibial plateau AC (b). *Gli1-LacZ-expressing* chondrocytes are significantly decreased in mutant AC (c-d). Tissue sections display patchy *Gli1-LacZ* positivity in the femoral condyle (c) and virtually no expression in the tibial plateau (d). (**E-H**) *Gli1-LacZ* at 8 weeks shows many positive cells in control AC of the femoral condyle (E) and tibial plateau (F). Tissue sections display similar patterning established in 3 week AC (e-f). *Gli1-lacZ-expressing* cells are still significantly reduced in mutant femoral condyle AC (G) but with maintained patchy expression from 3 weeks (g). In mutant tibial plateau AC, *Gli1-LacZ* is generally reduced in outer regions covered by meniscus and highly increased in the load-bearing center regions (H). Tissue sections from the load-bearing region reveal significantly increased *Gli1-LacZ* in all layers of cells (h). Scale bar = 100 µm.

### Hedgehog response patterning is coupled to tidemark patterning; both are disrupted in IFT88 mutants

Hedgehog signaling during embryonic development is tightly linked to tissue patterning through established gradients of ligand distribution. A notable example of this is neural tube formation where cells establish specific identities dependent on their exposure, and response, to hedgehog ligands (Ribes & Briscoe, 2009) and fine-tuned by a number of co-activators/antagonists (Allen et al., 2007, 2011; Holtz et al., 2013). Zonal chondrocyte patterning of the growth plate is also dependent on hedgehog exposure in a signaling loop that requires *Indian Hedgehog* (*Ihh*)-expressing pre-hypertophic chondrocytes and *Parathyroid related protein* (*Pthrp*)-expressing reserve zone chondrocytes (Kronenberg, 2006; St-Jacques et al., 1999). Hedgehog signaling patterns have not been well established in AC.

Using RNAscope, we assessed two major hedgehog response genes (*Gli1* and *Ptch1*) as well as *Ihh* in immature and mature AC and compared these patterns to the response when IFT88 was disrupted. Once again, we used confocal microscopy to image the FastRed detection and overlaid these results with DIC to visualize cell morphology, the tidemark (Fig. 8, dashed blue line), and the subchondral bone (sb, solid white line). First, we established patterning in immature AC at 3 weeks. In control AC, *Gli1* and *Ptch1* were expressed in overlapping domains surrounding the future tidemark in the first 3-4 cell layers in the femoral condyle (Fig. 8Aa-Ab) and 4-6 cell layers from the surface in the tibial plateau (Fig. 8Ba-Bb). Importantly, these data were consistent with those from our *Gli-LacZ* studies (Fig. 7A-B). In both regions only cells juxtaposed to the underlying subchondral bone expressed *Ihh* (Fig 8Ac and Bc). Two things were particularly notable about these patterns in immature AC: 1- the location of the future tidemark correlated strongly with *Gli1* and *Ptch1* expression, and 2- despite the *Ihh*-expressing cells being located always nearest the subchondral bone, the responding cells were closer to the surface in condylar cartilage than in the tibial plateau. In mutant AC at this stage, *Gli1* was significantly reduced in both femoral condyle and tibial plateau AC (Fig. 8Ad and Bd), consistent with *Gli1-LacZ* studies (Fig. 7C-D). *Ptch1* expression generally followed the same profile, but with some notable groups of positive cells in the femoral condyle (Fig. 8Ae, arrows and Be). Importantly, *Ihh* expression was unchanged compared to controls (Fig. 8Af and Bf). Thus, down regulation of signaling in this context is due to lack of response rather than lack of ligand expression.

**Figure 8.**
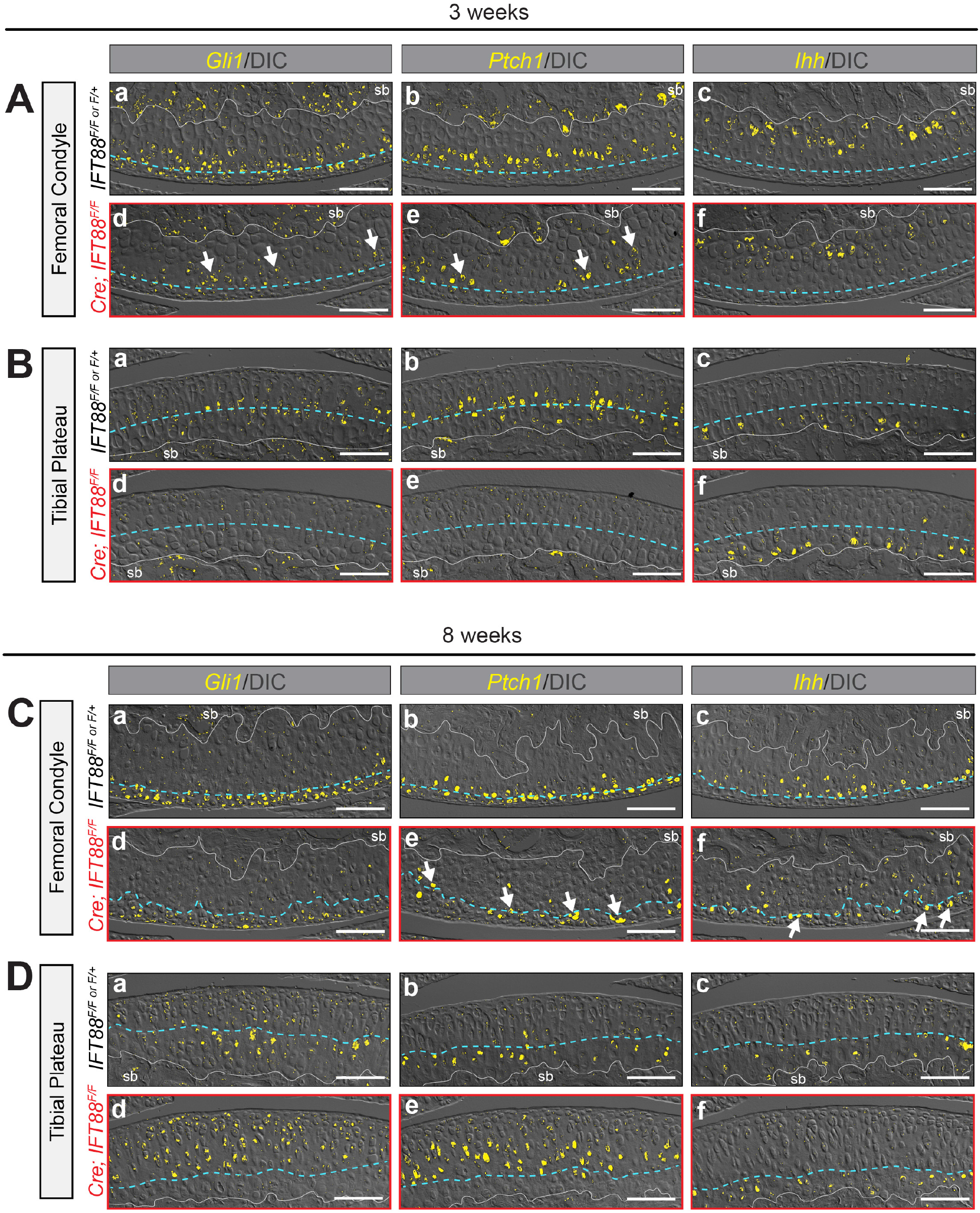
Hedgehog response patterning is coupled to tidemark patterning; both are disrupted in *IFT88* mutants. RNAscope detected by FastRed is visualized by confocal microscopy and overlaid with DIC imaging to visualize cell morphology and tidemark patterning. The blue dashed line in **A-B** represents the approximate level of the future tidemark in control AC and in **C-D** it follows the tidemark visualized by DIC. (**A**) Femoral condyle AC at 3 weeks. Control AC expresses *Gli1* in multiple cell layers near the AC surface (a), *Ptch1* in 1-2 cell layers around the presumptive tidemark (b) and *Ihh* in chondrocytes adjacent to the subchondral bone (c). Mutant AC displays decreased but patchy *Gli1* (d, arrows) and *Ptch1* (e, arrows) expression and normal *Ihh* in chondrocytes adjacent to the subchondral bone (f). (**B**) Tibial plateau AC at 3 weeks. Control AC expresses *Gli1* (a) and *Ptch1* (b) in similar layers surrounding the presumptive tidemark and *Ihh* in chondrocytes adjacent to the subchondral bone (c). Mutant AC displays decreased *Gli1* (d) and *Ptch1* (e) expression and normal *Ihh* in chondrocytes adjacent to the subchondral bone (f). (**C**) Femoral condyle AC at 8 weeks. Control AC expresses *Gli1* in multiple cell layers near the AC surface (a), *Ptch1* in 1-2 cell layers around the tidemark (b) and *Ihh* in chondrocytes just beneath the tidemark (c). Mutant AC displays decreased expression of *Gli1* (d), decreased and patchy expression of *Ptch1* that follows the irregular tidemark (e, arrows) and ectopic *Ihh* in non-mineralized chondrocytes (f, arrows). (**D**) Tibial plateau AC at 8 weeks. Control AC expresses *Gli1* (a) and *Ptch1* (b) in similar layers surrounding the tidemark and *Ihh* in chondrocytes just beneath the tidemark (c). Mutant AC displays increased and expanded expression of *Gli1* (d) and *Ptch1* (e) but normal *Ihh* just beneath the tidemark (f). scale bar = 100 µm.

We performed identical analyses in mature AC at 8 weeks of age. Femoral condyle AC of controls displayed *Gli1* expression in non-mineralized chondrocytes including surface zone cells (Fig. 8Ca), *Ptch1* expression in cells at the level of the tidemark (Fig. 8Cb), and *Ihh* expression in mineralized chondrocytes beneath the tidemark (Fig. 8Cc). Tibial plateau AC displayed overlapping *Gli1* and *Ptch1* in a zone 2-3 cell layers thick surrounding the tidemark (Fig. 8Da-b) and *Ihh* in mineralized chondrocytes beneath the tidemark (Fig. 8Dc). Femoral condyle cartilage of mutants displayed generally reduced *Gli1* above the tidemark (Fig. 8Cd) consistent with *Gli1*-*LacZ* studies. Interestingly, while *Ptch1* was also significantly reduced compared to control cartilage, the random distribution of persistent *Ptch1-*expressing chondrocytes correlated with the irregularly shaped tidemark (Fig. 8Ce, arrows). *Ihh* also persisted but was ectopically expressing in some non-mineralized chondrocytes (Fig. 8Cf, arrows). In tibial plateau cartilage, *Gli1* and *Ptch1* expression was highly increased in the load-bearing region (Fig. 8Dd-De) consistent with *Gli1-LacZ,* but unlike the condylar cartilage, *Ihh* remained exclusively expressed by non-mineralized chondrocytes beneath the tidemark (Fig.8Df). Taken together, these results provide evidence that hedgehog gradients coordinate tidemark patterning and that primary cilia guide zonal responses to provide unique tissue structure able to withstand local mechanical load.

## DISCUSSION

Our study of joint-specific loss of primary cilia function (IFT88-flox) provides new and probing insights into AC zonal patterning during postnatal development. We find that primary cilia are dispensable for embryonic joint formation, in line with previous *in vivo* studies (Chang et al., 2012; Irianto et al., 2014; Sheffield et al., 2018; Song et al., 2007; Yuan & Yang, 2015), but we have now uncovered the fact that they function primarily for AC maturation between adolescence (3 weeks) and adulthood (8 weeks). In this postnatal period, we discovered that the loss of primary cilia in AC reduced cell density and altered chondrocyte fate in either mineralized or non-mineralized cartilage, leading to disrupted ECM production/organization. Our most important and surprising finding was that loss of IFT88 alters tidemark patterning and topography in a manner mirroring alterations to hedgehog signaling. Through these analyses we were also able to establish, for the first time, the spatial patterning of *Ihh* responsiveness in mature AC.

Our study provides significant evidence that primary cilia and hedgehog signaling are key regulators of tidemark formation in mature AC. Ihh is required for joint specification during embryonic development (Koyama et al., 2007) and while the specific roles for hedgehog signaling in postnatal life have not been fully clarified genetic over activation of hedgehog signaling leads to early OA symptoms (Rockel et al., 2016). We show here that the emergence and location of the future tidemark in developing AC correlates strongly with patterning from Ihh signals. We show by RNAscope on tissue sections that even before the tidemark emerges at 3 weeks of age, hedgehog responding cells align with the future location of the tidemark in mature AC. Thus, our data establish a key causative link between patterning of mineralized and non-mineralized chondrocytes in AC and hedgehog signaling responses, guided by primary cilia. In future work, the mechanisms linking hedgehog signaling to tidemark patterning should be tested directly. Does the tidemark move dependent on a changing hedgehog response? What is the response in tissue integrity? Interestingly, tidemark advancement and duplication have also been associated with OA (Johnson, 1962; Lane & Bullough, 1980; Loeser et al., 2012; Oegema et al., 1997; Radin et al., 1991). Thus, these studies would delineate whether the movement in the tidemark advances the disease-state as hypothesized or serves another purpose yet to be discovered.

Our data also provide novel evidence that AC hedgehog signaling regulates *prg4* expression. *Prg4* (lubricin) is an important regulator of AC homeostasis (Das et al., 2019) and, in particular, prevents tissue and matrix degradation that would lead to OA (Chavez et al., 2019; Ruan et al., 2013). Understanding its transcriptional regulation may be key to preventing disease progression. Previous work has provided evidence of *prg4* regulation via TGFβ (Chavez et al., 2019), Creb5 (Zhang et al., 2021), Yap/Taz (Delve et al., 2020) and load-induced Cox2 (Abusara et al., 2013; Ogawa et al., 2014). In our present study, the decreased response to Ihh signaling detected in the non-mineralized chondrocytes of immature mutant AC was accompanied by reductions of *Prg4*. In mature AC *prg4/Ihh* were maintained at low levels in the femoral condyle, but expression of both genes increased dramatically in the load-bearing region of the tibial plateau. While most tissues follow the dogma that loss of the primary cilia leads to loss of hedghog signaling, the over-activation of hedgehog we see in our model is consistent with studies of embryonic limb development (Haycraft et al., 2005; Huangfu & Anderson, 2005) and postnatal musculoskeletal tissues (Chang et al., 2012; Kopinke et al., 2017) where Gli3 repressor plays a larger role in tissue function. Here, we provide evidence that this over-activation also has a mechanical loading component because only the load-bearing region of mature tibial plateau cartilage displays excessive hedgehog response. It is also plausible that reduced production of aggrecan and collagens might further compromise the response of chondrocytes to mechanical loading, and increase in *Prg4*. Studies designed to alter mechanical load in this model or alter hedgehog signaling separately would clarify the possible regulatory effect of hedgehog/load on *Prg4* expression.

In addition to the patterning deficits already acknowledged, we also found significant disruptions to the proteoglycan and collagen matrix in mutant animals. The overall mis-patterning of *Col2a1*- and *Acan*-expressing cells in mature AC can likely be attributed to hedgehog patterning disruptions that result in altered patterning of mineralized and non-mineralized chondrocytes in mature AC. However, it is notable that changes to ECM gene expression and production were not apparent in immature AC when hedgehog responsiveness was already altered. For this reason, we explored TGFβ and BMP signaling in order to gain further mechanistic insights into changes in ECM production in mature AC. We found that both signaling pathways were unchanged at 3 weeks but were significantly altered at 8 weeks, particularly in BMP signal response of the tibial plateau. Knowledge of the role(s) for BMP signaling in postnatal AC are limited, but two studies of conditional ablation of either *Bmpr1a* or *Bmp2* in embryonic joints (*Gdf5Cre*) reported no embryonic phenotype and a progressive loss of ECM during postnatal stages without hypertrophy or loss of Sox9 (Gamer et al., 2018; Rountree et al., 2004). These findings are strikingly similar to those we report here, particularly in the load-bearing region of the tibial plateau where we also observe a reduction of pSmad159 at 8 weeks of age. Possibilities for the loss of BMP responsiveness might include an altered adaptation to mechanical load without primary cilia or negative regulation by increased hedgehog signaling in the same location. To the latter point, hedgehog signaling is a known negative regulator of BMP during embryonic somitogenesis by enhancing the expression of BMP antagonists Noggin and Gremlin (Murtaugh et al., 1999; Stafford et al., 2011). Perhaps, a similar mechanism is at play here. In the femoral condyle, no such changes to BMP (or TGFβ) are noted and we also could not detect major changes to *Col2a1* or *Acan* gene expression. We believe that alterations to ECM in the femoral condyle are driven by cellular patterning defects that result in altered ECM organization while tibial plateau cartilage displays reduced ECM due to BMP/TGFβ dysregulation.

Our current study also raises new questions regarding the degree to which the primary cilium (and more specifically, IFT88) is required for mechanical adaptation in chondrocytes *in vivo. In vitro* studies of chondrocytes that lack primary cilia suggested a loss of mechanical transduction (Deren et al., 2016; Wann et al., 2012). Our present study shows a differential tissue response of hedgehog signaling in the femoral condyle and the load-bearing region of the tibial plateau. This altered response to ambulation load is indicative that tibial plateau chondrocytes are capable of “feeling” some mechanical load and have adapted without primary cilia. Underscoring this idea, recent work by Coveney et al. found that deletion of IFT88 using *AggrecanCreER* in adults led to thinner tibial plateau cartilage that was restored to normal thickness after giving animals free access to running wheels (Coveney et al., 2021). One possible mechanism for these adaptations could be through the pericellular matrix that also plays a major roles in articular chondrocyte mechanosensation (Guilak et al., 2006; Wilusz et al., 2012). Indeed, in our current study, mutant AC continues to display abundant perlecan localization in the pericellular matrix despite cellular mis-patterning. The data highlight the fact that cells are capable of mechanosensation through multiple mechanisms not necessarily requiring primary cilia, but that cilia may be required to coordinate for full adaptation.

On a broader level, our study underscores the differential topographical character of AC in normal joints and the differential phenotype due to the loss of primary cilia during development. Each joint must be uniquely fitted to the load-bearing and movement requirements of each anatomic location, and our study provides evidence that tidemark patterning is involved in this evolution. Specifically, we highlight differential tidemark patterning even in opposing AC in the same joint and a differential response to the loss of primary cilia. Femoral condyle AC showed more significant patterning disruptions, while tibial plateau AC showed more significant ECM loss. In this context, it is interesting to note that the mutant femoral condyle still elicited a significant reduction in mechanical stiffness. Our observations raise the important possibility that tissue patterning plays a much more significant role in overall AC integrity than currently appreciated. Taken into consideration and exploiting these facets of AC biology experimentally could lead to creation of more effective strategies for cartilage regeneration and repair.

## Material and Methods

### Animals

All studies involving mice were reviewed by our institutional IACUC at The Children’s Hospital of Philadelphia. All animals were handled, treated and cared for according to the approved protocols. *IFT88-flox* (Stock no. 022409) and *Rosa-TdTomato* mice (Stock no. 007914) were obtained from the Jackson Laboratory (Bar Harbor, ME, USA). *Gdf5Cre* mice (kindly provided by Dr. D. Kingsley) were described previously (Rountree et al., 2004) and line B was used in the present study. Wildtype littermate animals were used for controls.

### AFM-based nanoindentation

AFM-nanoindentation was applied to freshly dissected femoral condyle and tibial plateau cartilage from mice at 8 weeks of age (n = 5 animals), following the established procedure (Doyran et al., 2017; Han et al., 2019). Immediately after euthanasia, cartilage was dissected free of tendon and ligament tissues and glued onto AFM sample discs by a cyanoacrylate adhesive gel (Loctite 409, Henkel Corp., Rocky Hill, CT, USA). Throughout the procedure, tissues were immersed in PBS with protease inhibitors (Pierce 88266, Fisher Scientific, Rockford, IL, USA) to minimize post-mortem degradation. Nano-indentation was performed on the surfaces of cartilage on a Dimension Icon AFM (Bruker Nano, Santa Barbara, CA, USA) using a colloidal AFM probe. The probe was custom-made by attaching a polystyrene microsphere (R ≈ 12.5 µm, PolySciences, Warrington, PA) to a tipless cantilever (nominal k ≈ 5.4 N/m, cantilever C, HQ:NSC35/tipless/Cr–Au, Nano-AndMore, Watsonville, C, USA) using liquid epoxy glue (M-bond 610, SPI Supplies, West Chester, PA). For each sample, at least 10 random, different indentation locations were tested on each selected region of interest up to ∼1 µN force at 10 µm/s AFM z-piezo displacement rate. The effective indentation modulus, E_ind_, was calculated by fitting the entire loading portion of each indentation force delpth (F–D) curve to the Hertz model, where R is the tip radius and *ν* is the Poisson’s ratio (0.1 for cartilage (Buschmann et al., 1999)).

### Tissue Processing

Joints processed for frozen sectioning were injected with 4% PFA, fixed for 2 days at 4°C, decalcified in 20% EDTA for 6-10 days (depending on age), washed in PBS 3X15 minutes, equilibrated in 15% sucrose for 6-8 hours and in 30% sucrose for 16-36 hours (depending on age) and embedded in OCT. Sections were either take at 18μm thick for normal slide collection or 10 µm thick on Kawamoto’s cryofilm (Kawamoto & Kawamoto, 2021) to preserve tissue morphology. Joints processed for paraffin sectioning (basic histology and immunofluorescence) were injected with 10% NBF, fixed for 2 days at 4°C, decalcified in 20% EDTA for 6-10 days (depending on age), washed in PBS 3X15 minutes, washed in 70% ethanol 3X15min and processed to paraffin by automation overnight. Joints processed for RNAscope (Advanced Cell Diagnostics, Inc., San Jose, CA), were also processed for paraffin sectioning using Morse’s solution for decalcification as previously described (de Charleroy et al., 2021). All sections were cut at 6 µm thick and slides were prepared with mutant and littermate control sections on a single slide for each probe (N = 5 per time point).

### Histology/Immunofluorescence

Safranin O staining was performed using standard protocols and imaged using a Nikon Nikon Eclipse Ci-L upright microscope with attached DS-Fi3 camera (Nikon Corp., Tokyo, Japan). Immunofluorescence staining was carried out on frozen or paraffin sections depending on antibody optimization. For staining on frozen section, 1X PBS + 0.1% tween was used for all washes and incubations. Sections were blocked with 5% donkey serum and incubated overnight at 4°C against MMP13 (1:100, Abcam ab39012, Cambridge, UK) and for 2 hours at room temperature with a Cy3-conjugated secondary (1:500, Jackson Immunoresearch, West Grove, PA). For staining on paraffin sections antigen retrieval was performed by either 1mg/ml trypsin (Sigma, T7168, St. Louis, MO) or 10 μg/ml Proteinase K for 20 minutes at 37°C. For non-amplified staining, sections were blocked with 5% donkey serum (and Mouse-on-Mouse block as necessary from Vector Laboratories, Inc., Burlingame, CA) and incubated overnight at 4°C with an antibody against either Perlecan (1:50, Life Technologies, MA1-06821, Carlsbad, CA), Collagen2a1 (1:50, MilliporeSigma, MAB8887, Burlington, MA), CollagenX (1:50, MilliporeSigma, 234196), or acetylated tubulin (1:200, MilliporeSigma, MAB8887) and for 2 hours at room temperature with Cy3- or AF488-conjugated secondary antibodies (1:500, Jackson Immunoresearch). For amplified staining, all incubations after antigen retrieval were performed at room temperature. 1X PBS + 0.1% triton was used for permeabilization (20 minutes) and 1X PBS + 0.1% tween was used for all washes and buffers. Sections were treated with 3.6% H2O2 (made from 30% stock in water), blocked with 2.5-10% donkey serum (and Mouse-on-Mouse block as necessary from Vector Laboratories), incubated for 90 minutes with an antibody against either Sox9 (1:2000, Novus Biologicals, AF3075, Littleton, CO), Aggrecan (1:25, Developmental Studies Hybridoma Bank, 12/21/1-C-6, Iowa City, IA), pSmad2 (1:1500, Cell Signaling Technology, 3108S, Danvers, MA) or pSmad159 (1:800, Cell Signaling Technologies, 13820), for 1 hour with biotin-conjugated secondary antibodies (1:500, Jackson Immunoresearch), for 1 hour with Vectastain Elite ABC amplification kit (Vector Laboratories, Inc.) and developed for 10 minutes with fluorescein- or Cy3-conjugated Tyramide amplication system (Perkin Elmer, Shelton, CT). Amplified staining for Aggrecan was performed without detergent (1X PBS only) and included the following additional antigen retrieval steps prior to blocking: reduction with 10mM DTT for 2 hours at 37°C, alkylation with 40mM Iodoacetamide for 2 hours at room temperature, and 5 mg/ml Hyaluronidase (in 10% serum) for 1 hour at 37°C. All sections were counterstained with DAPI (1 mg/ml stock per manufacturer guidelines, Sigma, D9542) diluted 1:10,000 in wash buffer for 10 minutes and coverslipped using Prolong Gold (Invitrogen, P36930, Waltham, MA). A Leica DMi8CEL Advanced inverted SP8 confocal was used to collect fluorescent and DIC images (Leica Camera, Wetzlar, Germany). All stains were performed on at least 5 mutants and littermate controls per time point.

### Quantification from tissue sections

All quantification was carried out using ImageJ (National Institutes of Health, Software, Bethesda, MD) with FIJI plugins. All sections were chosen from the central region of the medial knee. For AC thickness measurements, Safranin O-stained sections were used. For each section (3-6 per animal), the thickest region of AC was identified and 8 lines were drawn manually and measured. An average and standard error was calculated from all lines in all sections per region of interest. For AC composition and cell density measurements DIC images overlaid with DAPI were used. The % composition was calculated by manually outlining and measuring the total cartilage, non-mineralized cartilage and mineralized cartilage area from at least 2 sections per animal. Cell density of the non-mineralized cartilage was calculated by counting (cell counter plugin) nuclei (DAPI in that area from at least 2 sections per animal. Density and %-positive cells for pSmad2 and pSmad159 were calculated from immunofluorescent-stained slides counterstained with DAPI and additionally imaged with DIC. DIC was used to outline the non-mineralized cartilage region per section (at least 2 sections per animal) and cells were counted using DAPI and pSmad staining.

### RNAscope

RNAscope *in situ* hybridization was carried out using RNAscope2.5 HD Detection reagent-RED (Advanced Cell Diagnostics, Newark, CA, USA) to visualize the spatio-temporal expression of *Adamts4* (cat. #497161), *Adamts5* (cat. #427621), *Col2a1* (cat. #407221), *Acan* (cat. #439101), *Prg4* (cat. #437661), *Gli1* (cat. #311001)*, Ptch1* (cat. #402811)*, Ihh* (cat. #413091). Briefly, sections were pretreated with a custom reagent and the probe was hybridized for 2 hours at 40°C in a custom oven. Signal was amplified with multiple reagents as per manufacturer’s protocols and final signal was detected and visualized by reaction with Fast Red substrate for 10-20 minutes (dependent on age) at room temperature. Companion sections were hybridized with positive (Cat No. 313911) or negative control probes (Cat. No. 310043) to assure signaling specificity. Sections imaged with brightfield were counterstained with hematoxylin, dried and sealed with Permount. Sections imaged with confocal microscopy were counterstained with Hoescht and cover slipped with Prolong Gold. A Nikon Eclipse Ci-L upright microscope with attached DS-Fi3 camera was used to capture brightfield images. A Leica DMi8CEL Advanced inverted SP8 confocal was used to collect fluorescent and DIC images.

### Statistical Methodologies

Data were analzed using Microsoft Excel and Prism 9 (Graphpad Software, Inc., San Diego, CA). Student’s *t* tests and one-way ANOVA was used to establish statistical significance with a threshold set at p < 0.05.

## Acknowledgements

This study was supported by NIH grants NIH/NIAMS F32AR074227 (to D.R.), R01AR062908 (to M.P.), R01AR074490 (to L.H.) and P30AR069619 (to Penn Center for Musculoskeletal Disorders). **Author Contributions:** D.R. and M.P. designed and managed the study. D.R., K.H., C.C., and E.K. performed animal experiments, data analyses, imaging, and quantifications. B.H. and L.H. performed AFM-nanoindentation and analysed data. D.R. wrote the manuscript with contributions and input from all authors. **Competing Interests:** The authors declare that they have no competing interests. **Data and material availability.** All data associated with this study are present in the paper and available from corresponding author upon reasonable request.

## SUPPLEMENTAL FIGURES

**Figure S1.**
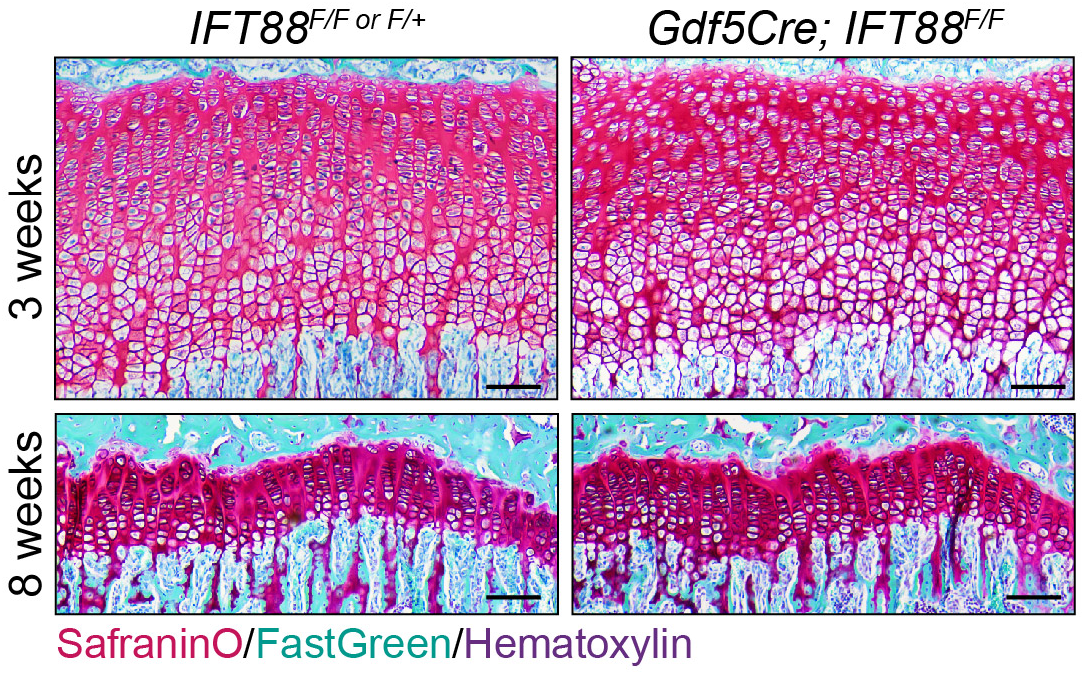
*Gdf5Cre/+; IFT88^F/F^* mutants show no alterations in growth plate patterning during postnatal development. Safranin O at 3 weeks and 8 weeks shows no change in growth plate morphology in joint-specific *IFT88* conditional mutants. scale bar = 100 µm.

**Figure S2.**
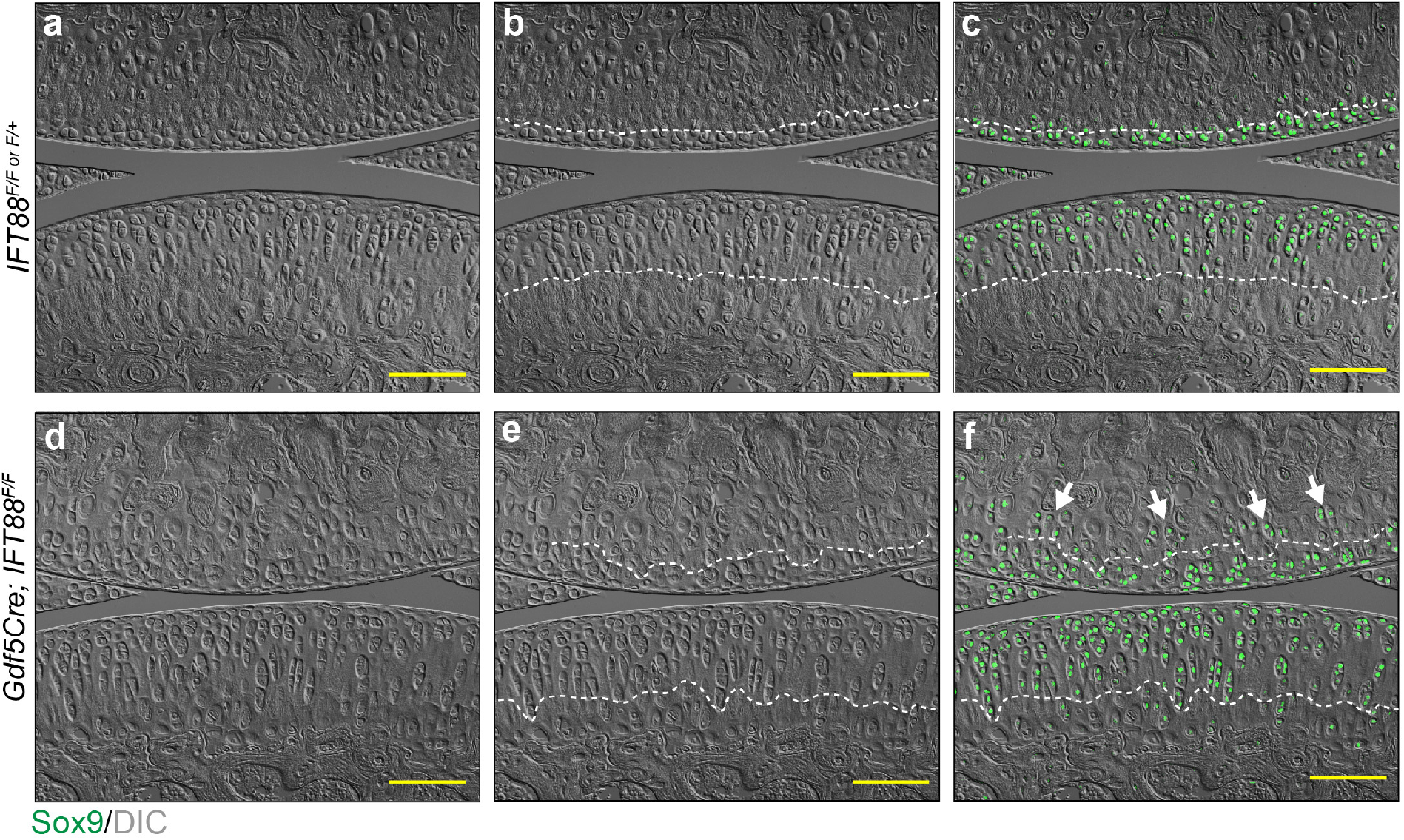
Sox9 overlaid with DIC-imaging to visualize the tidemark in controls and mutants. Sox9-stained images from Figure 3Dc-d. DIC imaging is shown without (a and d) and with (b and e) a white dashed line to denote the tidemark in controls and mutants and is overlaid with Sox9 (c and f) to show ectopic Sox9-positive chondrocytes in mutant femoral condyle AC (f, arrows). scale bar = 100 µm.

**Figure S3.**
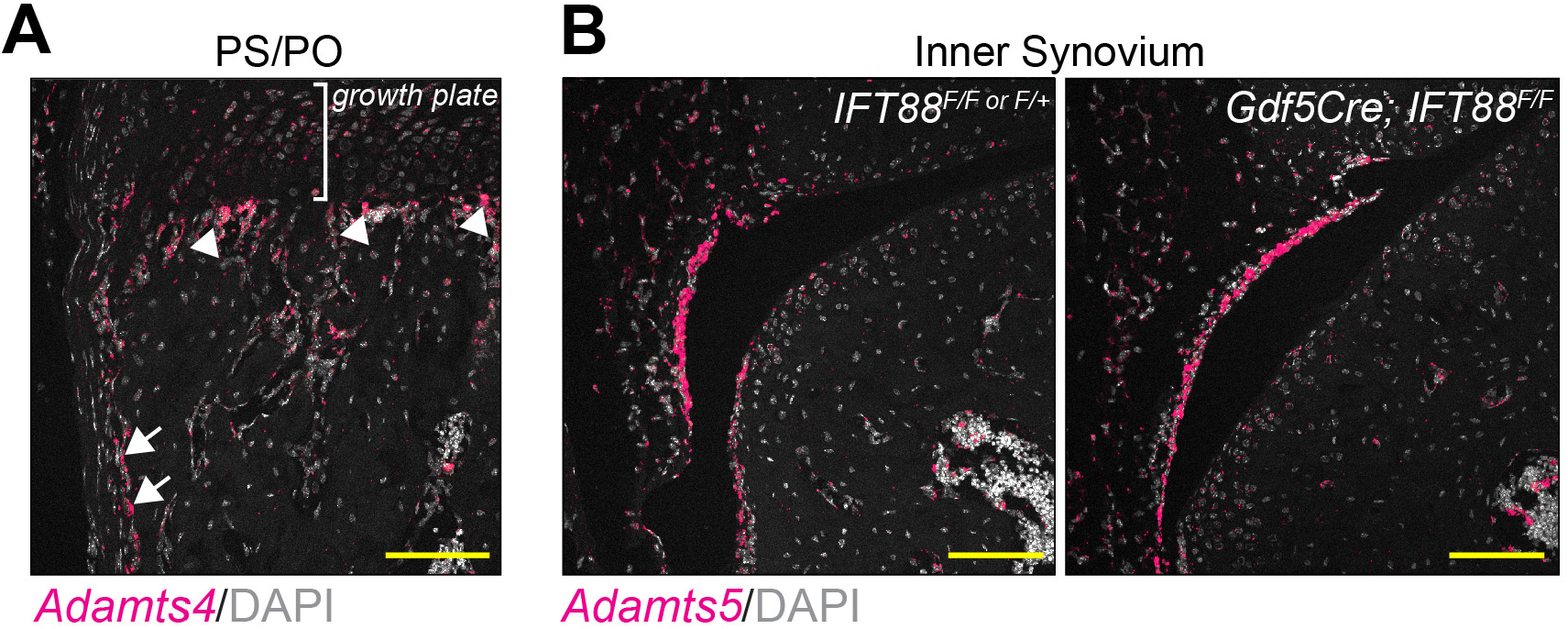
Expression of *Adamts4* and *Adamts5* outside of AC. RNAscope on 8 week joint/skeletal tissues developed with FastRed and imaged using confocal microscopy. (**A**) *Adamts4* is expressed the primary spongiosa beneath the growth plate (PS, arrowheads) and in the periosteum (PO, arrows). (**B**) *Adamts5* is expressed in the inner synovium of control and mutant joints. scale bar = 100 µm.

**Figure S4.**
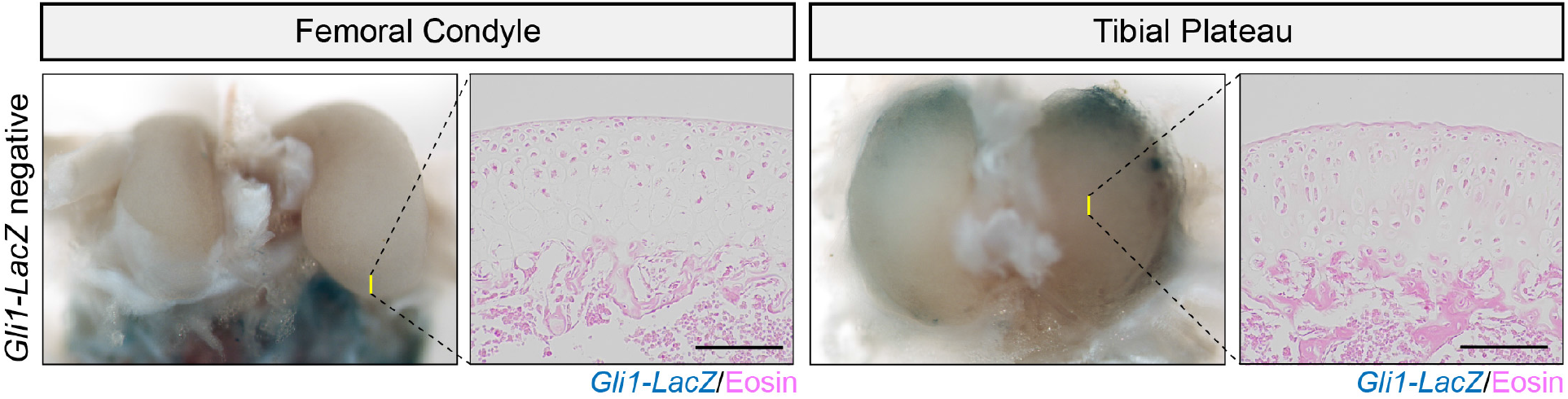
*Gli1-LacZ* negative tissues stained with β-gal. 3 week control, *Gli1-LacZ* negative AC stained with *Gli1-LacZ* positive AC from Figure 7. scale bar = 100 µm.

